# rareSurvival: rare variant association analysis for “time-to-event” outcomes

**DOI:** 10.1101/2021.12.19.473338

**Authors:** Hamzah Syed, Andrea L. Jorgensen, Andrew P. Morris

## Abstract

**Summary:** Rare variants have been proposed as contributing to the “missing heritability” of complex human traits. There has been much recent development of methodology to investigate association of complex traits with multiple rare variants within pre-defined “units” from sequence and array-based studies of the exome or genome. However, software for modelling time to event outcomes for rare variant associations has been under developed in comparison with binary and quantitative traits. We introduce a new command line application, rareSurvival, used for the analysis of rare variants with time to event outcomes. The program is compatible with high performance computing (HPC) clusters for batch processing. rareSurvival implements statistical methodology, which are a combination of widely used survival and gene-based analysis techniques such as the Cox proportional hazards model and the burden test. We introduce a novel piece of software that will be at the forefront of efforts to discover rare variants associated with a variety of complex diseases with survival endpoints.

**Availability & Implementation:** rareSurvival is implemented in C#, available on Linux, Windows and Mac OS X operating systems. It is freely available (GNU General Public License, version 3) to download from https://www.liverpool.ac.uk/translational-medicine/research/statistical-genetics/software/. Download Mono for Linux or Mac OS X to run software.

**Supplementary information:** Links to additional figures and tables are available at Bioinformatics online.

## Introduction

Methodology and software for the analysis of common variants within genome-wide association studies (GWAS) have been comprehensively developed and applied to a range of different endpoints, including survival data (Syed *et al*., 2017), resulting in the identification of thousands of loci for a wide range of complex traits and diseases. However, the common variant association signals in these loci account for only a small proportion of the genetic variance of the underlying trait. It is believed that rare genetic variants, defined as having minor allele frequency (MAF) less than 5%, may contribute to the “missing heritability” of human traits since they are not well covered through traditional GWAS, even after supplementation by high-density imputation. In the field of pharmacogenetics, in particular, there is evidence of a substantial contribution of rare coding variants to drug metabolism and transport (Gordon *et al*., 2014; Legge *et al*., 2016.).

Traditional methodologies for the analysis of GWAS lack power for rare variants (Moutsianas *et al*., 2015). Instead, multiple rare variants are most often jointly analysed within “functional units”, such as genes or pathways. Joint analyses are typically performed by assessing the association with rare variant “burden” or “dispersion” within the functional unit. These classes of tests each have their own benefits and limitations, dependent on the underlying genetic architecture of the trait under investigation. Moutsianas and Morris (2014) provide a comprehensive overview of available rare variant tests, and a discussion of architectures in which there is an advantage of using one method over another. The literature, to date, has focused on binary and quantitative traits, and is expanding rapidly with new methods becoming available. Software implementing a variety of binary and quantitative trait rare variant tests, such as EPACTS (http://genome.sph.umich.edu/wiki/EPACTS.), GRANVIL (Magi *et al.*, 2011) and VAT (Wang *et al*., 2014), are well developed to handle genome- or exome-wide data from sequencing or array genotyping. However, there has been very little activity regarding time to event phenotypes, which are particularly important within pharmacogenetics, where the outcome of interest could be time to disease remission or time to therapeutic drug dose.

Chen *et al.* (2014) have previously introduced the mathematical framework for rare variant associations with time to event outcomes for a general burden test and the dispersion-based sequence kernel association test (SKAT), focussing on a Cox proportional hazards model. They presented simulations to evaluate the relative power of burden tests and SKAT for time to event outcomes, coming to the same conclusions regarding the underlying genetic architecture as reported for binary and quantitative traits. This in turn provided us the framework to implement similar tests into freely available software. Software such as RVFam (Chen and Yang, 2015) has been developed for rare variant association analysis of family data. The software is designed to analyse survival traits using a Cox-proportional hazards model with shared frailty in each family by calling the coxph() function of the survival package in R. However, the software does not allow for more general survival models that allow for a deviation from the proportional hazards assumption.

To address this need, we have developed the rareSurvival software, which has been implemented using C# and run on Linux, Windows or Mac OS X operating systems. rareSurvival is capable of handling the scale and complexity of genome- or exome-wide data from sequencing or array genotyping..

## Features

### User interface

rareSurvival is a console application where users can either interactively use the software on a command line terminal or submit a script of commands to a high performance computing (HPC) capability. The program requires Mono (http://www.mono-project.com/) to run the software, because Linux and OS X are unable to compile C# code. A detailed workflow pipeline is presented in the supplementary information (Suppl. Fig. 1).

### Inputs

The user is required to specify three data files. The first must be a genotype file (.gen or .impute) or a variant call format (VCF) file, which contain genotype probabilities, dosages and/or hard genotype calls. The second must be a sample file that contains covariate and phenotype information for each individual. The file should contain a header that indicates the name of each column. This can be any text file extension. The third is the gene list file that contains information about genes on each row. The gene file can be in one of two different formats: (i) a .pos extension containing gene name, chromosome, base pair start position and base pair end position; or (ii) a .list extension containing gene name, chromosome and the variant identifier of each variant in the gene. The software can read in genotype files that are compressed, either in a .zip or .gz file extension. All files should have content separated by a single space or tab.

The user is required to specify additional information, including the outcome of interest (i.e. survival time and censoring indicator) and any covariates to be adjusted for in association analyse.. Furthermore, a specification of the range of genomic regions to be analysed in the gene list file (to enable parallel processing), the choice of survival analysis method and the rare variant test, the MAF and info score thresholds and the name of the results output file. The default values for MAF and info score are 0.05 and 0.9, respectively. Suppl. Table 1 contains full description of commands, along with a command line and script example.

### Validation algorithm

rareSurvival will convert the genotype probabilities into a variant coded under an additive dosage model. If hard genotype calls or dosages are present in the file then these are directly used without conversion. The hard genotype calls from VCF files can be in the form 0/0, 0/1, 1/1 or 0|0, 0|1, 1|1.

Each SNP in turn is matched with gene location or variant list in the gene list file, along with the MAF and info score threshold. Errors occur if the user has specified an incorrect command or states a variable name that cannot be found in the header of data files. rareSurvival will terminate the task and it will need re-submission. The program also handles missing values (coded as “NA”) within the sample file. If a subject has missing values for survival time or a covariate specified in the model, they removed from the analysis.

### Output

The output from the analysis will be saved in a text file with name specified by the user. The file contains the gene name and/or covariate name, chromosome number, total number of rare variants in region, total number of variants, sum of MAF’s, mean MAF, sum of info score, mean info score, coefficient value, hazard ratio, standard error, confidence intervals (only for Cox proportional hazards), Wald test p-value, likelihood ratio test p-value, model likelihood ratio test p-value .

## Methods

The software currently supports analysis using a burden test (with optional Madsen-Browning weighting) within a Cox proportional hazards or Weibull regression model framework. The burden score across variants within a gene is introduced into the Cox proportional hazards model or Weibull regression model as a variable. Coefficients are then derived along with the corresponding p-value using a Wald or Likelihood ratio test.

## Results

To exemplify the efficiency of the software, we conducted simulations based on Illumina Exome Array genotype data obtained from 2,120 individuals from two cohorts of elderly Swedish individuals (Lithell *et al.*, 2000). We simulated phenotype information using SurvivalGWAS_Power (Syed *et al.*, 2016) based on two randomly selected genes containing our causal variants. Variants were assigned to genes based on base pair position using the gene list file which links coding variants (defined by a strict annotation) to genes. Analysis was completed on a total of 73952 SNPs over 12432 genomic regions across 23 chromosomes for 2120 individuals. For a detailed summary of the simulation setup see supplementary information Table 1.2. For results refer to Supplementary information Figure 2

The complete analysis was run using 5 computer nodes (40 cores). Each node consisted of a HP Proliant DL170h G6 server, 2 Intel Xeon(R) E5520 2.27GHz quad-core CPUs, 36 GB memory and 1 TB of local storage. Running the gene-based analysis using a burden test in a Cox proportional hazards model with no additional covariates took 5.4033 hours to complete. Supplementary information table 1.3 has a breakdown of computational runtime dependent on number of covariates.

## Discussion

We have introduced a novel analysis tool for rare variant association studies with time to event outcomes. rareSurvival implements burden tests of association within a regression framework, allowing for Cox proportional hazards and Weibull models.

After identifying the genomic regions in our simulation study for which our causal variants were located, the next step would be to pinpoint which variants within the gene are leading this signal of association. This is a crucial next step, with a need for further method development similar to Lin *et al*. (2016) and application into this software.

With the increasing availability of sequence data, particularly in the field of pharmacogenetics, and the pursuit of analysing more complex time to event outcomes, future versions of rareSurvival will incorporate multifaceted survival models together with the latest rare variant analysis methods, to account for non-proportional hazards, competing risks, variable variant effect sizes and directions.

In conclusion, rareSurvival will play a key role in the discovery of novel genes associated with patient response to treatment for a range of complex human diseases, eventually leading to the personalisation of therapeutic intervention. Using rareSurvival for the combination of sequence based genetic data and detailed clinical assessments will help us better understand the genetic architecture of complex diseases, facilitating translation of statistical results into practical solutions to advance disease prediction and treatment.

Conflict of Interest: none declared.

