## Supplementary material for "rareSurvival: rare variant association analysis for “time-to-event” outcomes": Supplimentary Information

Supplementary Information

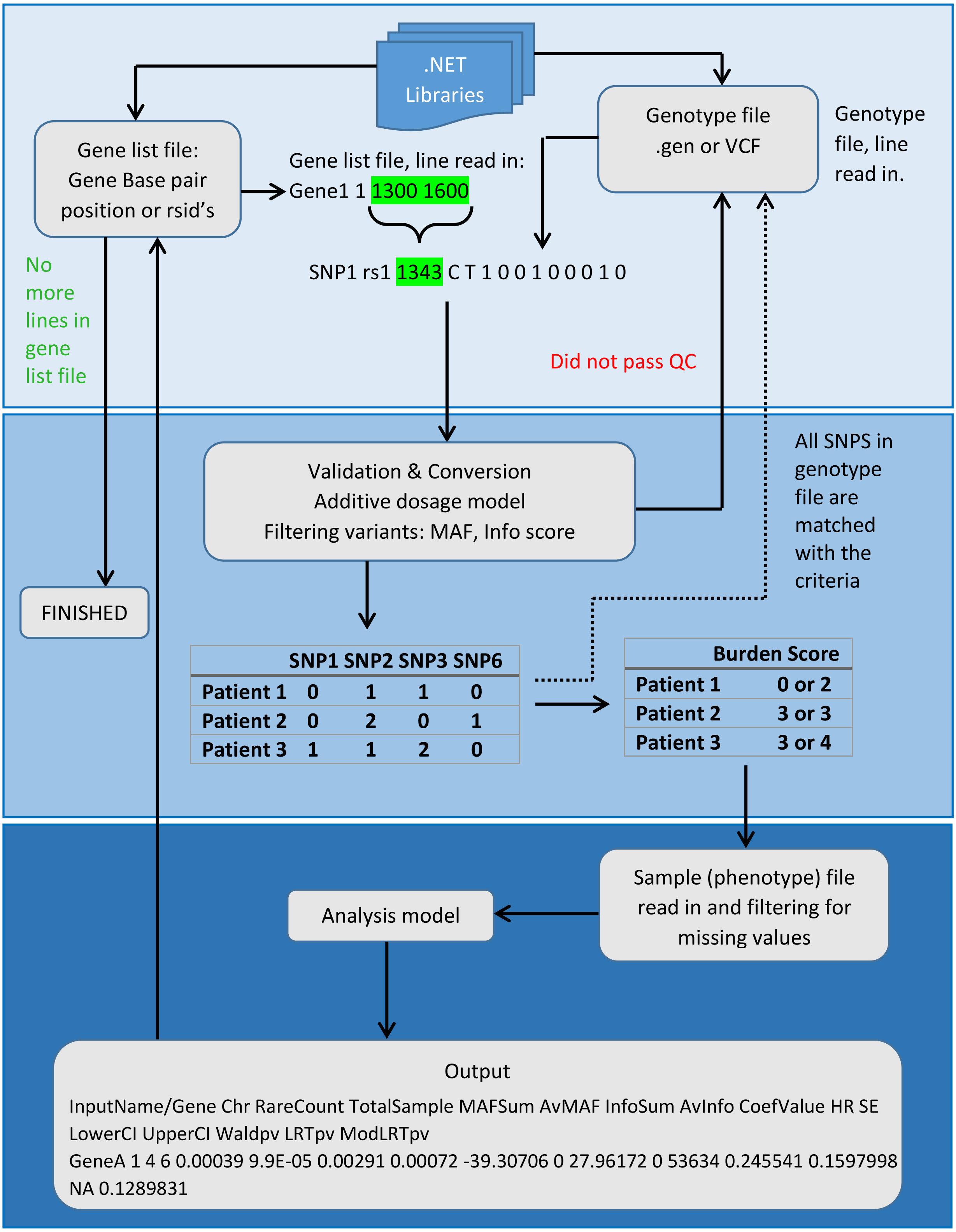


Fig. SD1.1. Workflow example of rareSurvival, from data input to results output. Scenario:

| Command | Description |
|---|---|
| **-gf=** | This specifies the genotype file. Typically .gen, .vcf, .impute, .gen.gz. |
| **-sf=** | This specifies the sample file (.sample, .txt). |
| **-mf=** | This specifies the gene list file. |
| **-threads=** | Number of threads. On a multi-core system, multiple threads can execute tasks in parallel, with each core executing a single or multiple threads. |
| **-t=** | This specifies the time to event (column heading name) in the sample file. |
| **-c=** | This specifies the censoring indicator/outcome in the sample file. |
| **-cov=** | This specifies the covariates to adjust for in the model. Each one separated by a comma (,). Categorical factors need to be converted to binary as software only assumes continuous or binary covariates. |
| **-lstart=** | This specifies the line in the gene list file at which the start position of analysis will occur. Used to break large files into small batches for parallel computing. |
| **-lstop=** | This specifies the line in the gene list file at which the end position of analysis will occur. Typically the number of lines is equal to the number of genomic regions in the file. |
| **-chr=** | This specifies the chromosome number to be output in the text file. |
| **-p=** | Enter “onlygene” if only the results from the gene analysis are to be output. <optional> |
| **-m=** | This specifies the choice of survival analysis method. This is either “cox” for the Cox proportional hazards model or “weibull” for the parametric Weibull regression model. |
| **-rm=** | Specifies the choice of rare variant analysis method. This is “burden” for the BT with unit weighting or “mbweight” for the Madsen & Browning weighted BT. |
| **-maf=** | Specifies the MAF threshold for inclusion of SNPs in the analysis. |
| **-info=** | Specifies the info score (imputation quality) threshold for inclusion of SNPs in the analysis. |
| **-o=** | This specifies the name of the file for output to be saved in. e.g. name.txt |
| **-help** | Outputs a full list of commands and usage help. |

Table. SD1.1. List of commands available in the software and their corresponding usage description.

**Command line example**

Assuming all data files and software are in the same folder, the command line in a Linux terminal for the analysis of 100 genes on chromosome 6 with 1 additional covariate using a BT in a Cox proportional hazards model is as follows:

```
mono raresurvival.exe -threads=4 -gf=data.vcf -sf=data.sample –
mf=list.txt -t=event_times -c=outcome -cov=covariate1 -chr=6 -
lstart=0 -lstop=100 –maf=0.01 -info=0.9 -m=cox -rm=burden -
p=onlygene -o=output.txt
```

This line is also filtering for SNP’s with MAF less than equal to 0.01. We are analysing the data on a 4 core computer therefore we have specified 4 threads to spread the analysis output over. All 4 threads will analyse a different chunk of the data and output to the same file. Each command is separated by a space. The user can specify the exact location of the data files and where the output file will be saved. e.g. `/DIRECTORY/DATA/output.txt`

An example of a shell script (.sh) to distribute the analyses between 10 computer cores within a Linux cluster, using a sun grid engine batch system is as follows:

```
#!/bin/bash
#$ -o stdout
#$ -e stderr

DIRECTORY=/rareSurvival #Location of software and data
str1=0 #Start position in gene list file
str=100 #Number of genes/lines in gene list file
no_of_jobs=10 #Number of cores
inc=`expr \( $str - $str1 \) \/ $no_of_jobs` #Increment

#SGE_TASK_ID takes values 1:no_of_jobs
nstart=`expr \( $SGE_TASK_ID - 1 \) \* $inc’
nstop=`expr $nstart + $inc – 1`
mono $DIRECTORY/raresurvival.exe -threads=4 -gf=$DIRECTORY/data.vcf
-sf=$DIRECTORY/data.sample –mf=$DIRECTORY/list.txt -t=event_times -
c=outcome -cov=covariate1 -chr=6 -lstart=0 -lstop=100 –maf=0.01 –
info=0.9 -m=cox –rm=burden -p=onlygene -
o=$DIRECTORY/output${SGE_TASK_ID}.txt
```

To submit the script file you can use the command:

```
qsub -t 1:10 script.sh
```

If your cluster does not have the ***qsub*** command installed and you have access to all available cores from the head node, submit using the following:

```
sh script.sh
```

The first gene was *FNDC1* (chromosome 6), which contains 25 rare variants. We randomly selected 15 variants to be causal, with the same direction of effect, and simulated survival times for each individual. The second gene was *OR5B17* (chromosome 11), which contains 4 rare variants. We simulated survival times based on all 4 causal variants but with varying low to moderate effect sizes with the same direction of effects.

| Gene | Unique variant identity number | Reference /alternate allele | Position | MAF | Function | Simulated effect size |
|---|---|---|---|---|---|---|
| FNDC1 | - | C/T | 159636159 | 0.00023585 | Unknown | 0.6 |
| | rs186515442 | G/A | 159646577 | 0.00023585 | Missense | 0.6 |
| | rs200758408 | G/A | 159659662 | 0.00165094 | Missense | 0.6 |
| | rs61746218 | C/T | 159667972 | 0.00259434 | Missense | 0.6 |
| | rs200171920 | A/T | 159672511 | 0.00023585 | Missense | 0.6 |
| | rs180849332 | C/T | 159687181 | 0.00165094 | Missense | 0.6 |
| | - | G/A | 159692377 | 0.00283019 | Unknown | 0.6 |
| | rs200925962 | A/G | 159692428 | 0.0009434 | Missense | 0.6 |
| | rs202080149 | A/G | 159644604 | 0.00023585 | Missense | 0.6 |
| | rs202114028 | G/A | 159653261 | 0.00023585 | Missense | 0.6 |
| | rs199900169 | G/A | 159636039 | 0.0009434 | Missense | 0.6 |
| | rs201387402 | A/G | 159644575 | 0.0004717 | Missense | 0.6 |
| | - | A/G | 159653612 | 0.0004717 | Unknown | 0.6 |
| | rs7763726 | A/G | 159670100 | 0.02004717 | Missense | 0.6 |
| | rs186422799 | G/A | 159653921 | 0.00683962 | Missense | 0.6 |

| | | | | | | |
|---|---|---|---|---|---|---|
| OR5B17 | rs144440324 | G/C | 58125740 | 0.000235849 | Missense | 0.1 |
| | rs55810057 | A/G | 58125774 | 0.018632075 | Missense | 0.15 |
| | rs199650837 | A/T | 58126153 | 0.000943396 | Stop-gained | 0.2 |
| | rs140465731 | C/T | 58126340 | 0.000471698 | Missense | 0.05 |

Table. SD 1.4. List of causal variants used in simulation study. Based on Exome array.
**Stop-gained**: leading to a gain of a stop codon.
**Missense**: leading to an amino acid change.

**Simulation model:** List of causal rare variants. We simulated survival times for each individual, $i$, using a proportional hazards model incorporating each selected causal variant within the scale parameter of a Weibull distribution. The probability density function of the Weibull distribution $t_i = Weibull(1, 50 \times e^{\sum_{j=1}^{m} \delta_j G_{ij}})$
$j$ represents the number of causal rare variants.

$$\delta_j = \begin{cases} 0.6, FNDC1 \\ (0.1, 0.15, 0.2, 0.05), OR5B17 \end{cases}, m = \begin{cases} 15, \ FNDC1 \\ 4, \ OR5B17 \end{cases}$$

Censoring was randomly simulated using an exponential distribution with scale parameter 10.
Number of censored observations = …

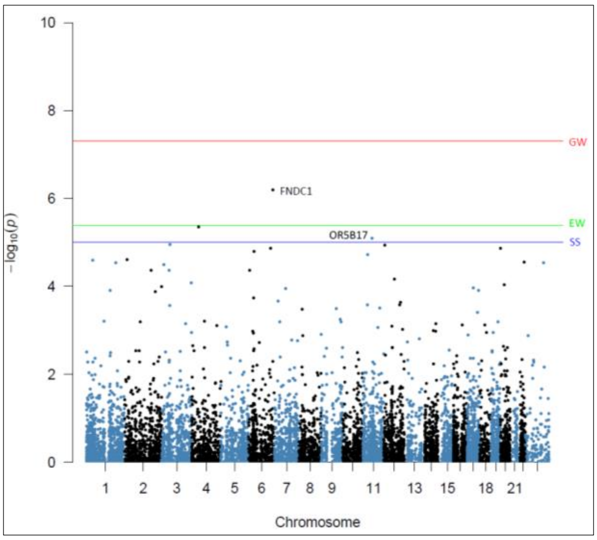


Fig. SD1.2. Manhattan plot of results from Swedish cohort exome chip data using a Burden test in a Cox proportional hazards model, for the identification of genes with causal variants. X-Axis is chromosome number and Y-Axis is Wald test -log10(p-value). Each blue and black point represents a gene.

Supplementary Figure 2 presents the results of application of rareSurvival under a Cox proportional hazards model. The first causal gene, ***FNDC1,*** is associated at exome-wide significance (Bonferroni correction for 12,432 genes, $p < 4.0 \times 10^{-6}$) with the simulated time to event outcome ($p=6.4 \times 10^{-7}$, 0.000943, 3.80E-05,1,0.04). The second causal gene, ***OR5B17,*** shows strong association, but not at exome-wide significance, which is expected as it contains fewer causal variants with low effect sizes ($p=8.1 \times 10^{-6}$, Sum of MAF = 0.000472, Average MAF=0.000118, InfoSum = 1, InfoAV=0.25).

| Number of additional covariates | Runtime of complete analysis for simulation study setting. From data input to results output. |
|---|---|
| 0 | 324.198 minutes |
| Treatment (Binary) | 378 minutes |

| | |
|---|---|
| Treatment (Binary) & Age (20-50) | 445.2 minutes |

Table. SD1.3. Computational runtime of software with and without additional covariates.

Treatment was simulated using a Binomial distribution. Age was simulated with a Uniform distribution.

The runtimes in table.SD1.3 are highly dependent on computing resources available to the user, size and compression of genotype file and number of individuals.
